# A commercial antibody to the human condensin II subunit NCAPH2 cross-reacts with a SWI/SNF complex component

**DOI:** 10.1101/2020.11.07.372599

**Authors:** Erin E. Cutts, Gillian C Taylor, Mercedes Pardo, Lu Yu, Jimi C Wills, Jyoti S. Choudhary, Alessandro Vannini, Andrew J Wood

**Affiliations:** Division of Structural Biology, The Institute of Cancer Research, London SW7 3RP, United Kingdom; MRC Human Genetics Unit, Institute of Genetics and Molecular Medicine, The University of Edinburgh, Edinburgh, EH4 2XU, UK; Cancer Research UK Edinburgh Centre, Institute of Genetics and Molecular Medicine, University of Edinburgh, Edinburgh, EH4 2XU, UK

## Abstract

Condensin complexes compact and disentangle chromosomes in preparation for cell division. Commercially available antibodies raised against condensin subunits have been widely used to characterise their cellular interactome. Here we have assessed the specificity of a polyclonal antibody (Bethyl A302-276A) that is commonly used as a probe for NCAPH2, the kleisin subunit of condensin II, in mammalian cells. We find that, in addition to its intended target, this antibody cross-reacts with one or more components of the SWI/SNF family of chromatin remodelling complexes in an NCAPH2-independent manner. This cross-reactivity with an abundant chromatin-associated factor is likely to affect the interpretation of protein and chromatin immunoprecipitation experiments that make use of this antibody probe.

## Main Text

In order to identify proteins that physically interact with the condensin II complex, we performed affinity purification mass spectrometry using a polyclonal antibody raised against the C-terminus of the NCAPH2 subunit (Bethyl Labs A302-276A, referred to hereafter as B276). Immunoprecipitation was performed from mouse embryonic stem cell (mESC) whole cell extract, without crosslinking, using either B276 or a rabbit IgG control. The abundance of co-purified proteins was then assessed by Label Free Quantification. As expected, all 5 subunits of the pentameric condensin II complex were significantly enriched in the Ncaph2 immunoprecipitate compared to the IgG control (Figure 1A). In addition, numerous components of the SWI/SNF complex family were also enriched to a similar or greater degree than known condensin II subunits. Indeed, the most significantly enriched protein identified in these experiments was the Smarcc1/Baf155 subunit.

**Figure 1:**
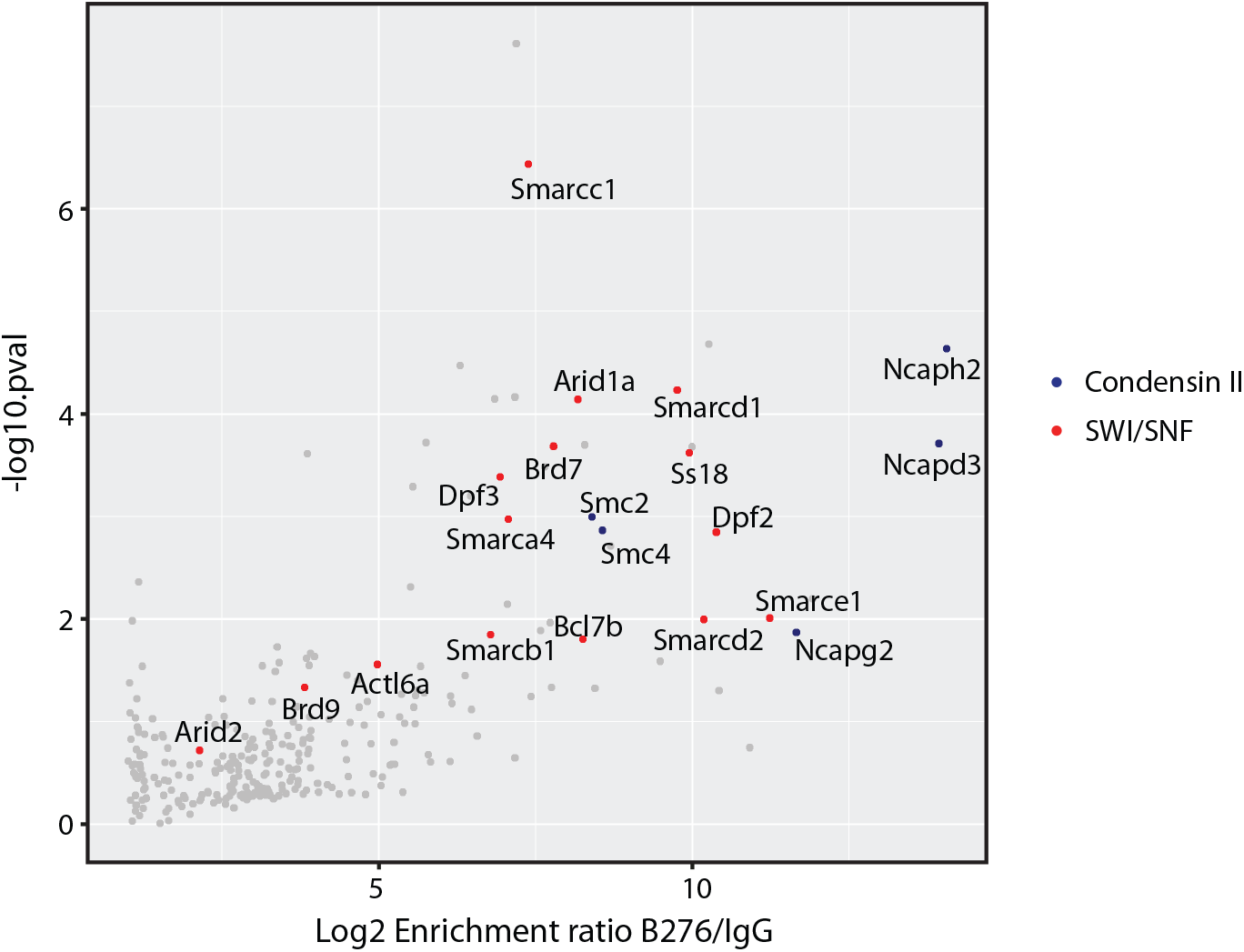
Co-immunoprecipitation of Condensin II and SWI/SNF complex subunits by a commercial anti-NCAPH2 antibody. Scatter plot showing the enrichment of proteins in immunoprecipitations conducted using the Bethyl Labs anti-NCAPH2 (B276) antibody versus rabbit IgG control (n = 4, from cultures grown and processed in parallel), as determined by mass spectrometry and label-free quantification. The x-axis depicts mean Log2 fold-enrichment. The y-axis shows the Log_10_ probability of enrichment from Bayes-moderated t-tests. Condensin II and SWI/SNF subunits are labelled in blue and red, respectively.

In an independent experiment performed in a different laboratory, immunoprecipitations were performed using nuclear lysates from human HEK-293T cells, either after crosslinking with formaldehyde or with non-crosslinked material. Co-purified proteins were identified using MS/MS and analysed with the SAINTexpress algorithm (Teo et al. 2014) to determine fold enrichment scores (Table 1 and 2). For both crosslinked and non-crosslinked cells, all components of the condensin II pentamer were identified with high enrichment scores, as well as numerous components of the SWI/SNF complex, confirming the independent results observed in mESC.

**Table 1:** Enrichment of Condensin II and SWI/SNF in IP using crosslinked nuclear lysate with B276.

| Hit Number | Uniprot ID | GENE | Enrichment Score (FC_A) | Anti-CAP-H2 Peptide Count, run 1 | Anti-CAP-H2 Peptide Count, run 2 | IG Peptide Count, run 1 | IG Peptide Count, run 2 |
| --- | --- | --- | --- | --- | --- | --- | --- |
| 1 | Q6IBW4 | NCAPH2 | 6.72 | 114 | 89 | 0 | 0 |
| 2 | O95347 | SMC2 | 5.91 | 104 | 68 | 0 | 0 |
| 3 | Q9NTJ3 | SMC4 | 5.85 | 97 | 75 | 0 | 0 |
| 4 | Q92922 | SMARCC1 | 4.86 | 72 | 67 | 0 | 0 |
| 5 | P42695 | NCAPD3 | 4.52 | 73 | 51 | 0 | 0 |
| 6 | Q86XI2 | NCAPG2 | 4.3 | 61 | 58 | 0 | 0 |
| 7 | P51532 | SMARCA4 | 4.14 | 51 | 65 | 0 | 0 |
| 9 | O14497 | ARID1A | 3.2 | 34 | 48 | 0 | 0 |
| 15 | Q969G3 | SMARCE1 | 2.62 | 28 | 31 | 0 | 0 |
| 24 | Q8NFD5 | ARID1B | 2.42 | 17 | 38 | 0 | 0 |

**Table 2.**
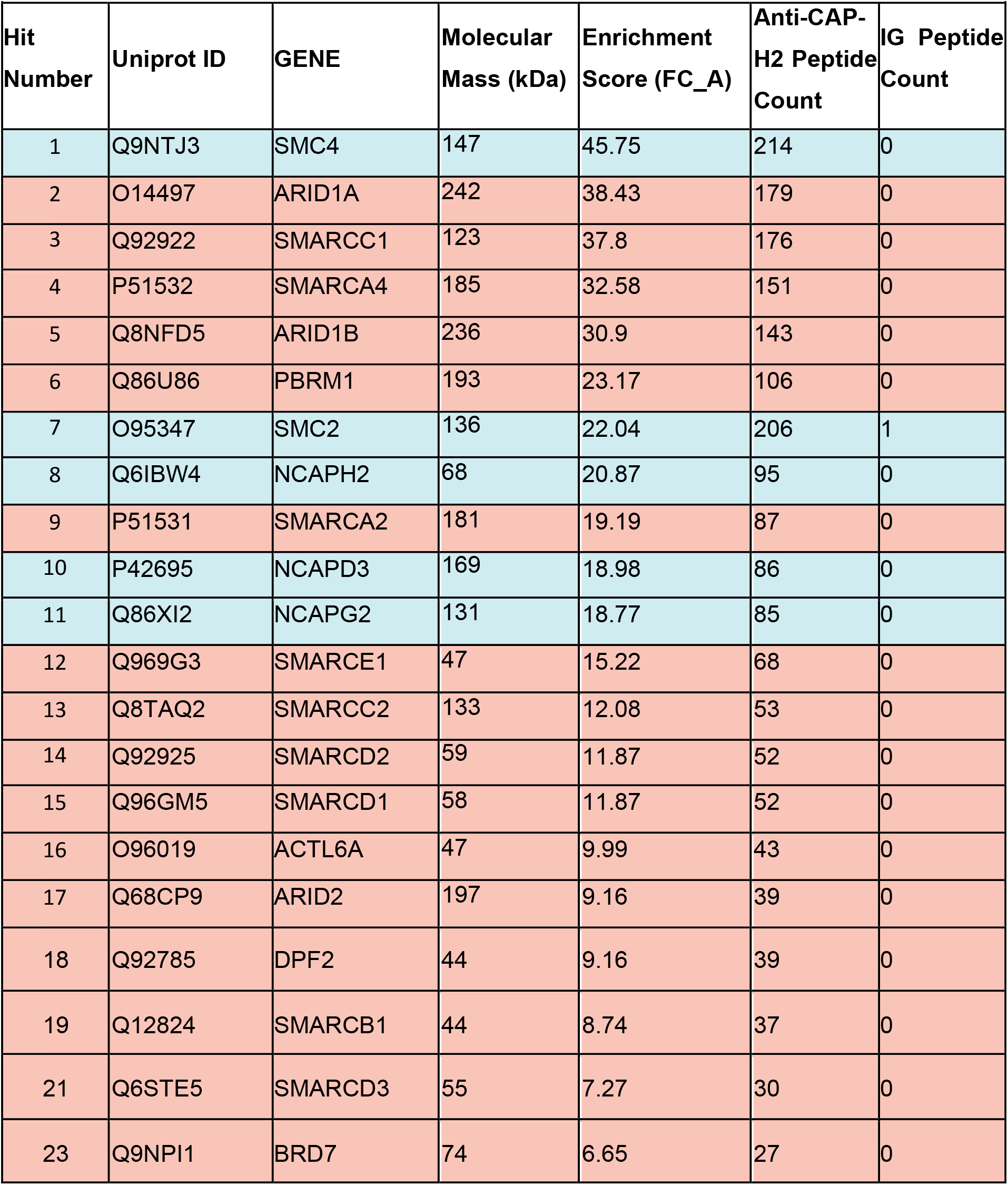
Enrichment of Condensin II and SWI/SNF in IP using non-crosslinked nuclear lysate with B276.

These results were unexpected, because previous biochemical characterisation of SWI/SNF complexes did not identify condensin subunits as robust interactors (Ho et al. 2009; Kadoch et al. 2013). We therefore considered two possibilities: a direct physiological interaction between condensin II and SWI/SNF, leading to *bona fide* co-purification, or, on the contrary, a direct recognition of one or more SWI/SNF components by the B276 anti-NCAPH2 antibody, resulting in an experimental artefact.

To distinguish these possibilities, we first performed reciprocal immunoprecipitations using B276 (Fig. 2A) and an antibody against the SWI/SNF subunit Smarca4 (Fig. 2B). Confirming the Mass Spectrometric analysis, B276 immunoprecipitated both the Smc4 subunit of condensin and the SWI/SNF subunit Smarca4 (Fig. 2A). However in the reciprocal experiment, the anti-Smarca4 antibody robustly immunoprecipitated the SWI/SNF component Arid1a, as expected, but Ncaph2 was not detected by western blots using the B276 probe (Fig. 2B).

**Figure 2:**
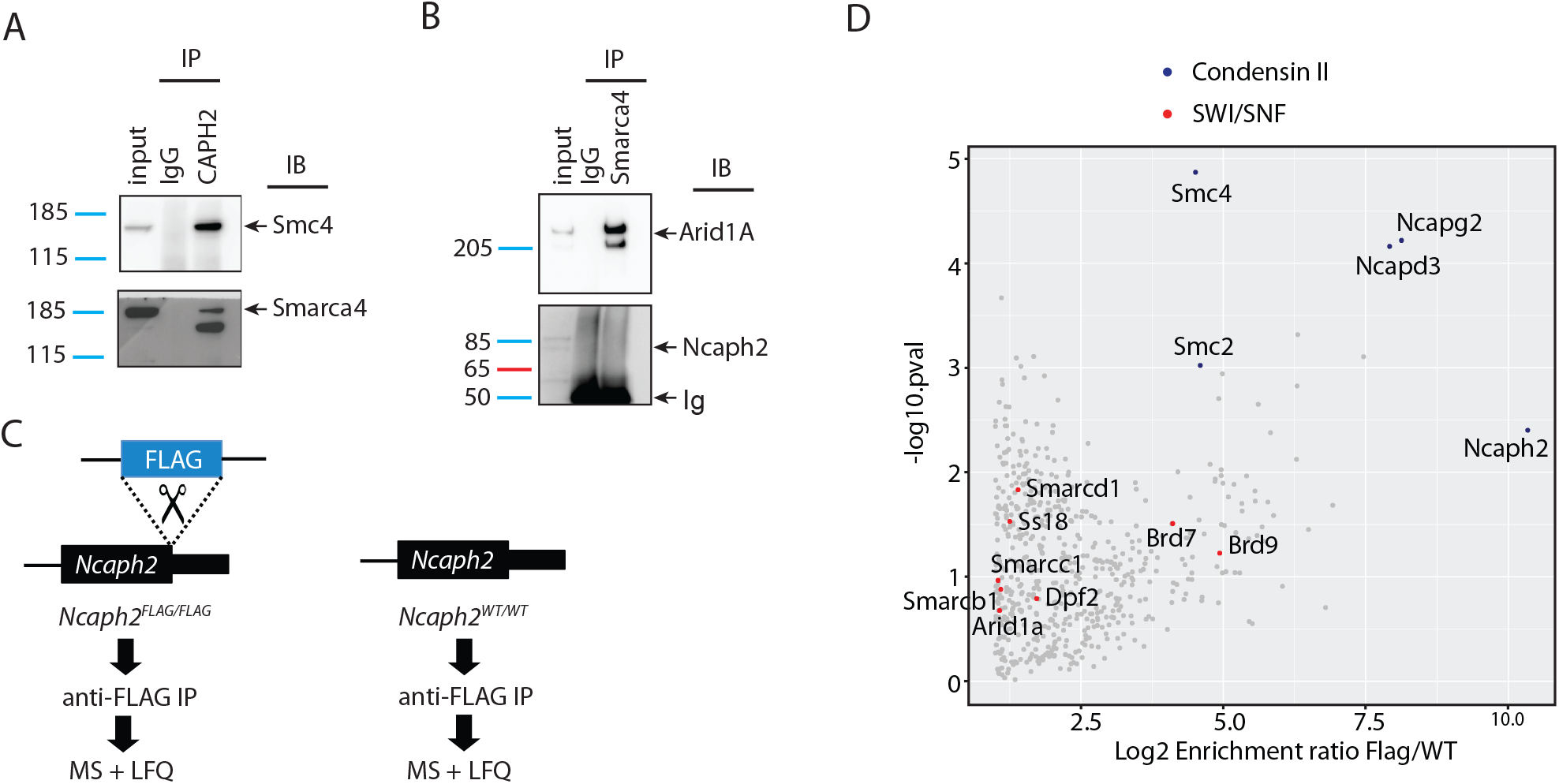
Independent strategies do not support physical interactions between condensin II and SWI/SNF subunits. **A**. Immunoblots (IB) show enrichment of the condensin subunit Smc4 and the SWI/SNF subunit Smarca4 in immunoprecipitations (IP) conducted from thymic whole cell extract using the B276 antibody compared to IgG control. **B**. Immunoblots show enrichment of the SWI/SNF subunit Arid1a (∼270 KDa), but not the condensin II subunit Ncaph2 (∼80 KDa), in immunoprecipitations conducted using an anti-Smarca4 antibody compared to IgG control. **C**. Schematic illustration of anti-FLAG affinity purification mass spectrometry strategy using NCAPH2FLAG/FLAG mouse ESCs and untagged control. **D**. Scatter plot showing protein enrichment in the immunoprecipitation experiments shown in panel C (n = 4, from cultures grown and processed in parallel). The x-axis depicts mean Log2 fold-enrichment in Ncaph2FLAG/FLAG (Flag) versus untagged control cells (WT). The y-axis shows the Log_10_ probability of enrichment from Bayes-moderated t-tests. Condensin II and SWI/SNF subunits are labelled in blue and red, respectively.

In a second independent experimental setup, CRISPR-mediated homology-directed repair was used to homozygously integrate a cassette encoding a double FLAG tag at the 3’ terminus of the endogenous *Ncaph2* open reading frame in mESCs (*Ncaph2*^*FLAG/FLAG*^, Fig. 2C). Seamless integration of the epitope tag was confirmed by Sanger sequencing. Anti-FLAG antibodies were then used to perform affinity purification from whole cell extract of *Ncaph2*^*FLAG/FLAG*^ or control *Ncaph2*^*+/+*^ mESCs, without crosslinking, and peptides were identified by mass spectrometry and label free quantification as described above. All 5 condensin II subunits were significantly enriched in the *Ncaph2*^*FLAG/FLAG*^ immunoprecipitate, whereas SWI/SNF subunits were not (Fig. 2D). Thus, although several SWI/SNF subunits were immunoprecipitated by the B276 anti-NCAPH2 antibody in both human and mouse cells, we were unable to validate these interactions using two independent approaches. This suggested that the immunoprecipitation of SWI/SNF by B276 occurred independently of the NCAPH2 target in both mouse and human cells.

To directly test this hypothesis, we made use of an HCT-116 Human cell line in which C-terminally tagged NCAPH2 protein can be rapidly depleted by the addition of indole-3-acetic acid (auxin) to the culture medium (Fig. 3A)(Takagi et al. 2018). Whole cell extracts were prepared from cells cultured either in the presence or absence of auxin treatment (5 hours), and immunoprecipitations were conducted using B276, or using a custom anti-NCAPH2 antibody raised in our laboratory (Methods).

**Figure 3:**
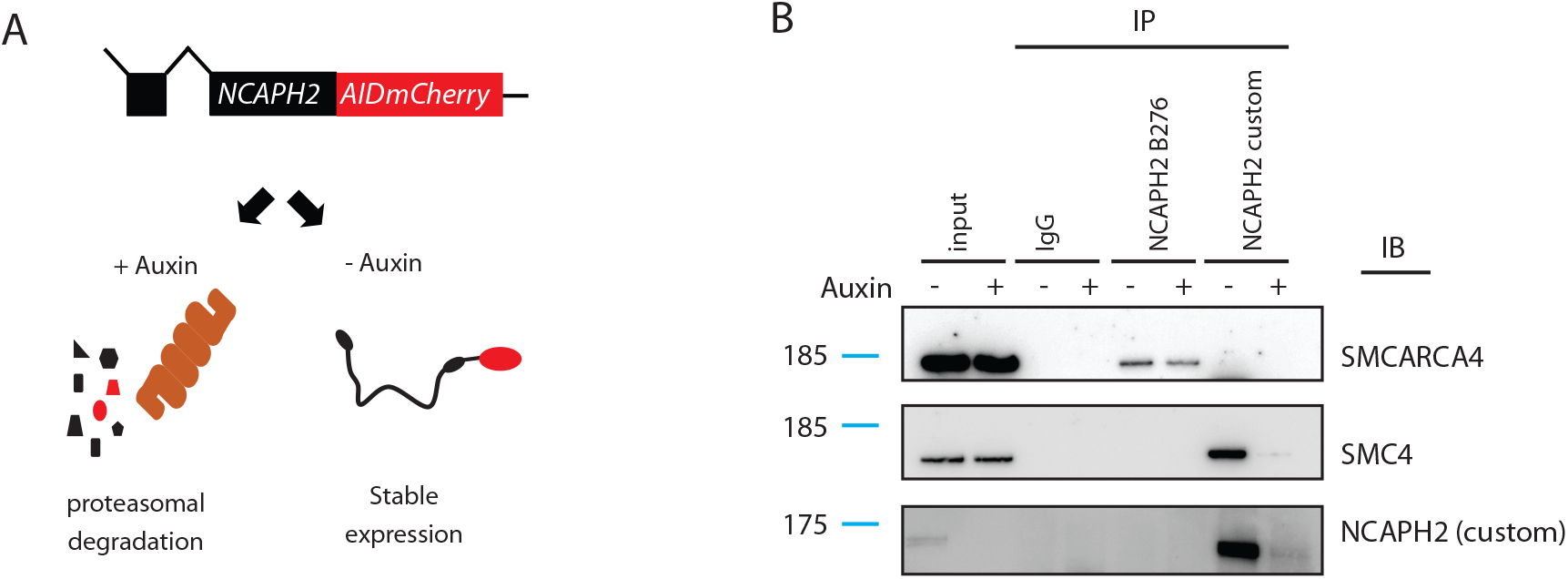
Immunoprecipitation of SWI/SNF by B276 does not require Ncaph2. A. Schematic illustration of the auxin inducible degron system used to rapidly deplete NCAPH2 protein from HCT-116 cells. B. Immunoblots (IB) conducted on whole cell extract immunoprecipitated (IP) using the indicated antibodies. IPs were conducted in the presence (+) versus absence (-) of auxin-mediated NCAPH2 depletion.

In the absence of auxin treatment the custom antibody efficiently immunoprecipitated both NCAPH2 and the condensin subunit SMC4, but this was prevented by the auxin-mediated depletion of NCAPH2 (Fig. 3B). The SMARCA4 subunit of SWI/SNF was not detectable by immunoblot in either condition. In contrast, the B276 antibody immunoprecipitated SMARCA4 but not NCAPH2 or SMC4 in this cell line (Fig. 3B). Why B276 can immunoprecipitate Ncaph2 from mESC and HEK-293T cell extracts but not from this HCT-116 cell line is unclear, but could be attributable to the C-terminal AID:mCherry tag in the latter. Importantly, the immunoprecipitation of SMARCA4 by B276 was unaffected by auxin-mediated depletion of NCAPH2. These results confirm that the immunoprecipitation of SWI/SNF complexes by B276 occurs independently of the intended NCAPH2 bait, and therefore do not reflect *bona fide* physical interactions between condensin II and SWI/SNF.

To further investigate details of SWI/SNF components binding to the 276 antibody, a western blot was performed on HEK 293T cell nuclear lysate as well as a recombinant human condensin II complex (Kong et al. 2020)(Figure 4A). While recombinant protein resulted in one band which ran in the position of NCAPH2 at around ∼80 kDa, the blot of nuclear lysate produced two bands, one at around 80 kDa, likely corresponding to NCAPH2, and an additional band around 135 kDa. We then performed an IP using B276 followed by western blotting with either B276 or an antibody recognising SMARCC1 whose result suggested that both NCAPH2 and SMARCC1 were pulled down. However, the additional 135 kDa band was observed at the same respective height in both blots developed either with B276 or with an anti-SMARCC1 antibody (Figure 4B). This finding leads us to suspect that B276 binds with considerable affinity to SMARCC1, in addition to NCAPH2, accounting for the considerable enrichment of SMARCC1 in the B276 IP mass-spectrometry data (Figure 1A, Tables 1 & 2).

**Figure 4:**
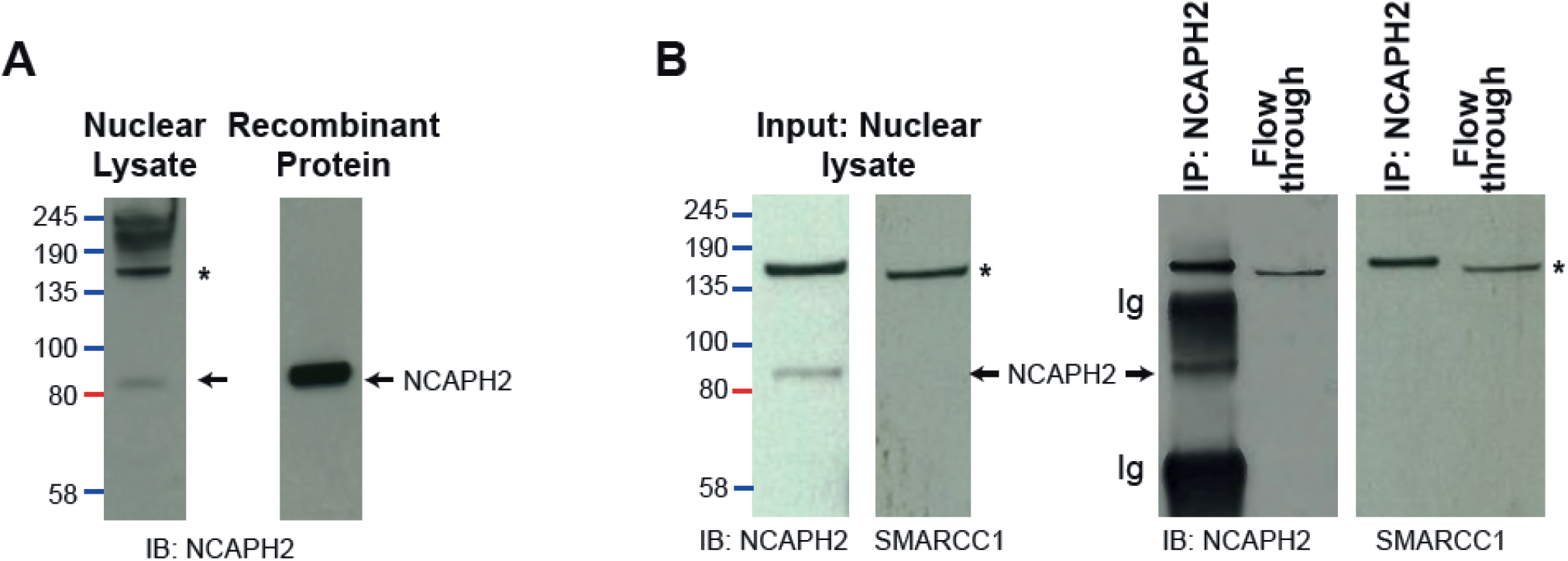
B276 has affinity to a 135 kDa protein which runs at the same position as SMARCC1. **A**. Immunoblot (IB) of nuclear lysate from formaldehyde crosslinked 293T cells probed with B276 yields two bands, a specific band at ∼80kDa, that is also present when B276 is used to probe a blot prepared from recombinant condensin II complex (Kong et al, 2020), and a non-specific band (*) at ∼135kDa. **B**. Immunoprecipitation (IP) performed with B276 using 293T cell nuclear lysate, followed by immunoblot probed with either B276 or mouse Anti-SMARCC1 (sc-32763), demonstrating that the non-specific band in B276 (*) runs in the same position as SMARCC1.

The cross-reactivity of the B276 antibody with another protein, specifically an abundant chromatin-associated factor, may affect the interpretation of protein and chromatin immunoprecipitation experiments performed using this antibody (Wu et al. 2019; Li et al. 2015). In particular the notion that Condensin II and components of the SWI/SNF chromatin remodeller complexes colocalize on a genome-wide scale (Wu et al. 2019) should be revisited in light of the presented results.

## Materials and Methods

The experiments presented in this paper were performed independently between two research laboratories, without prior coordination. To reflect the fact that different protocols were followed in each laboratory we provide their methods separately, indicating the relevant Figure panels in each case.

### Experiments performed at the MRC Human Genetics Unit

#### Cell culture

E14 mouse embryonic stem cells (Figures 1 & 2) were cultured in GMEM (Gibco) supplemented with 15% FCS, 0.1mM β-mercaptoethanol, penicillin (10,000 units/mL) and streptomycin (650µg/mL), 0.3mg/mL L-Glutamine, 1000 units/mL human recombinant LIF, 1mM sodium pyruvate and non-essential amino acids (Sigma), at 37°C, 5% CO_2_, and 3% O_2_.

HCT-116 NCAPH2-AID-mCherry cells (Figure 3) were cultured in DMEM (Gibco) with 10% (v/v) foetal calf serum (FCS), penicillin (10,000 units/mL) and streptomycin (650µg/mL) at 37°C, 5% CO_2_, and 3% O_2_.

#### Genome Editing

To generate the NCAPH2^FLAG/FLAG^ mESC line used in Figure 2, the following sgRNA protospacer sequences were cloned into pX335-U6-Chimeric_BB-CBh-hSpCas9n(D10A):

guide 1 = GGTGGAAAGTAGTATATACC

guide 4 = ACTCAAGGCTGGGCCATGGA

The homology-directed repair template was ordered as a double stranded DNA fragment (gBlock, IDT) and cloned using MluI sites into pEGFP-C1 (Clontech).

Repair template sequence:

ACGCGTAAGTGGCCTGGGACACAAGGGATGGGGCAGCGGCCCTGGACTCTACTGACACCTTGTTTCCACAGGCTAATGACTACACAGTGGAGATCACTCAGCAGCCAGGACTGGAGGCAGCTGTGGACACAATGTCTCTGAGACTGCTCACACACCAGCGAGCCCACACCCGCTTCCAGACCTATGCTGCACCATCCATGGCCCAGCCTGACTACAAGGACGACGATGACAAGGACTACAAGG ACGACGATGACAAGTGAGTGGACAGCACTGAGGCAGGGGTGGAAAGTAGTATATACCTGGAGGTCTTTGCCCCTAATGTGCTATGGGGCCATTCACTCCAGTGCTGCCTCCTGGCTGGCCTAGCCTAATAAAGTGTTGCTACCCCACCTGTTCACCGGACAGACTATTTAAATGAGCTGCTGGTACAGAGCACATGCACAGATTAAGTACATCCATTTAATGACAGGGCCTAGGCAATAGGTATAGTTCAGACGCGT

The pX335 derivative plasmids encoding sgRNAs and Cas9 nickase were transfected in combination with the repair template into E14 mESC using Lipofectamine 2000 according to manufacturer instructions. Transfected cells were FACS-purified based on GFP fluorescence and seeded at limiting dilution, then individual colonies were picked, expanded, screened by PCR band-shift assays, and then homozygous integration was verified by Sanger sequencing.

### Immunoprecipitation Mass Spectrometry

For experiments presented in Figures 1 and 2, trypsinised suspensions of E14 mESC (WT or NCAPH2-FLAG) were lysed in 1% triton buffer (20mM Tris-HCl, 137mM NaCl, 1% triton X-100, 2mM EDTA, protease inhibitors (Calbiochem), phosphatase inhibitors (Roche)). Protein concentration was quantified using a Pierce BCA Protein Assay Kit (Thermo, 23228) following manufacturer’s instructions. 1mg of total protein was used for immunoprecipitation, which was performed on a Kingfisher Duo robot using 1µg of antibody, magnetic beads, and 500µl of lysate at 2mg/mL. Lysate, antibodies and beads were incubated with mixing at 5 °C for 2 hours, then beads were transferred for two washes in lysis buffer without detergent, and three washes in TBS. Following tryptic digest (2M urea, 100 mM tris, 1 mM DTT, 5µg/mL trypsin), peptides were separated and ionised on Ultimate 3000 series Nano UHPLC using a C18 RP packed emitter in a column oven (IonOpticks, Sonation, Proxeon). Ionised peptides were analysed on a Q Exactive Plus (Thermo) in data-dependent mode and searched against the 2018_01 release of Uniprot mouse reference proteome using MaxQuant version 1.6.1.0, with LFQ (Cox et al. 2014). Statistical analysis was conducted using Wasim Aftab’s implementation of Kammers et al 2015 (https://github.com/wasimaftab; (Kammers et al. 2015)).

### Immunoprecipitation for Western Blot

For immunoprecipitations in Figure 2A & 2B, single cell suspensions of thymus tissue were generated by gentle dissociation of whole thymus tissue through 40 µm filters and lysed in 1% triton buffer as previously. For immunoprecipitations in Figure 3B, HCT-116 NCAPH2-AID-mCherry cells (Takagi et al. 2018) (+/-5 hour treatment with 500µM auxin), were typsinised and lysed in 1% triton buffer. Protein concentration was determined by BCA assay as described above. 1mg total protein lysate was incubated with 30µg Protein A Dynabeads (Invitrogen 10001D) and 1µg antibody (anti-NCAPH2, bethyl labs, anti-NCAPH2 custom, anti-SMARCA4/Brg1, Santa-Cruz, or anti-rabbit IgG) at 4°C overnight. Beads were washed 4 times rotating at 4°C for 5 minutes in IP wash buffer (10mM Tris-HCl, 2mM EDTA, 1% tween 20, 0.5% Triton X-1000) and once in TE buffer (10mM Tris-HCl, 1mM EDTA). Beads were then resuspended in 40uL 1x NuPage LDS sample buffer with 1x NuPage Sample reducing buffer (ThermoFisher NP0007, NP0004) before denaturation via boiling at 95 °C for 5 minutes. Samples were used immediately or stored at −20 °C.

### Western Blots

For western blots in Figure 2A and 2B, denatured protein lysates (10µL/sample) were loaded on to NuPAGE 4-12% Bis-Tris or 3-8% Tris-Acetate 1.0 mm Mini Protein Gels (Invitrogen, NP0321, EA0375PK2) buffer alongside PageRuler Protein Ladder (5 µL/lane; Thermo Scientific, 26616) and run in pre-chilled 1X MOPS Buffer (Thermo, NP0001) or NuPage Tris-Acetate SDS running buffer (Invitrogen LA0041). Samples were typically run at 100 Volts for 90 minutes. Transfers were performed onto Immobilon-P PVDF membranes (Millipore, IPVH00010) via wet transfer; PVDF membranes were pre-soaked in 100 % methanol (Fisher, 10284580) and rinsed briefly in Transfer Buffer (25 mM Tris (AnalaR, 103156X), 200 mM glycine (Fisher, G-0800-60), 20% methanol, 0.02% SDS (IGMM Technical Services)). Genie Blotter transfer device (Idea Scientific) was assembled with the gel and PVDF membrane placed between two layers of cellulose filter paper (Whatman, 3030-917) inside the loading tray. Once the apparatus was prepared, Transfer Buffer was filled to the top of the Genie Blotter and transfer proceeded for 90 minutes at 12 volts. Samples were blocked with 5% milk powder (Marvel) in TBS (10mM Tris-HCl, 150mM NaCl) plus 0.1% Tween20, with constant agitation, for 1 hour at room temperature.

Primary antibodies (detailed in Table 3) were added to the block solution at the dilution shown in the antibody Table. Membranes were incubated in the antibody dilutions with constant agitation, at 4 °C overnight. Membranes were washed in TBS-Tween 20 solution (0.1% Tween20; 3 washes x 10 minutes). HRP-conjugated secondary antibodies were also diluted in the block solution (with 0.1% Tween20), and membranes were incubated with secondary antibody dilutions under constant agitation at room temperature for 1 hour. Membranes were then washed in TBS-Tween20 solutions (0.1% Tween20, 3 washes x 10 minutes). Membranes were stained with SuperSignal West Pico PLUS HRP substrate (Thermo Scientific) and then imaged with an ImageQuant LAS 4000 (GE Healthcare).

### Experiments performed at the Institute of Cancer Research

#### Immunoprecipitation on HEK293T cells

For the experiments presented in Tables 1 & 2, HEK 293T cells were seeded at 5×10^6 cells and grown to 70-90% confluence in 10 cm plates in DMEM (Gibco) with 10% (v/v) foetal calf serum (FCS), penicillin (10,000 units/mL) and streptomycin (650µg/mL) at 37°C, 5% CO_2_. Non-crosslinked cells were washed 3 times in with PBS, flash frozen and stored at −80°C. Fixed cells were crosslinked with 1% formaldehyde for 10 minutes, then quenched with 0.125 M of glycine for 5 minutes, before washing three times with PBS and freezing.

Each immunoprecipitation was performed using cells from ten 10cm plates, using 100 µL of Protein A Dyna beads (Invitrogen) precoated with 10 µg of Rabbit anti-NcapH2 (A302-276A, lot 1) or rabbit IG (CST #2729) and washed with PBS +0.01% tween. Cells were lysed with Lysis buffer (10 mM HEPES pH 8.0, 85mM KCl, 0.5% NP-40, protease and phosphatase inhibitor (Pierce)) for 10 minutes at 4°C, then spun at 500 g to pellet nuclei.

Immunoprecipitation using crosslinked cells was performed twice, in two independent experiments. Crosslinked cells nuclear lysate was prepared by lysing nuclei using RIPA buffer (50 mM Tris pH 8, 150 mM NaCl, 1% NP-40, 0.1% SDS, 0.5% DOC, protease and phosphatase inhibitors, 1mM DTT) and a Dounce homogeniser. DNA was fragmented with sonication, and DNA fragment size was confirmed to be less than 500 bp by running a sample on 1.5% agarose, stained with SYBR safe (Thermo Fisher). DNA was further digested by incubating with Benzonase with 2 mM MgCl_2_ for 60 minutes, before being spun at 12,000 g. Nuclear lysate was incubated with dyna beads for 2 hrs, and beads were washed 5 times with RIPA buffer.

Un-crosslinked cell immunoprecipitation was performed once, as described in Bode et al (Bode et al. 2016). Briefly, nuclear lysate was prepared by solubilising nuclear pellet with nuclear lysis buffer (50 mM HEPES (pH7.9), 5mM MgCl_2_, 0.2% TritonX-100, 20% glycerol, 300 mM NaCl, and protease and phosphatase inhibitor) using a Dounce homogeniser with a tight pestle and incubating with Benzonase (Sigma) for 60 minutes at 4°C before being spun at 12,000 g. Nuclear lysate was diluted 1 in 2 with nuclei lysis buffer without NaCl before incubating with Dyna beads for 2 hrs at 4°C. Beads were washed 5 times with nuclear lysis buffer with 150mM NaCl.

After IP for both crosslinked and un-crosslinked samples, on bead digestion was performed as described in Hiller et al (Hillier et al. 2019). Briefly, beads were washed into 50 mM Ammonium Bicarbonate (FLUKA-40867) and digested with 0.1ug/ul of Trypsin (ROCHE-11047841001) overnight at 37°C. Peptides were collected, filtered using a Millipore Multiscreen HTS plate, lyophilised and resuspended in 20 mM TCEP (SIGMA-646547) with 0.5% formic acid (FISHER-A117-50) for analysis with mass-spectrometry.

### Mass spectrometry

The LC-MS/MS analysis was on the Orbitrap Fusion Tribrid or Fusion Lumos mass spectrometer coupled with U3000 RSLCnano UHPLC system. All instrument and columns used were from Thermo Fisher. The peptides were first loaded to a PepMap C18 trap (100 µm i.d. x 20 mm, 100 Å, 5 µm) at 10 µl/min with 0.1% FA/H2O, and then separated on a PepMap C18 column (75 µm i.d. x 500 mm, 100 Å, 2 µm) at 300 nl/min and a linear gradient of 4-32% ACN/0.1% FA in 90 min with the cycle at 120 min. Briefly, the Orbitrap full MS survey scan was m/z 375 – 1500 with the resolution 120,000 at m/z 200, with AGC (Automatic Gain Control) set at 40,000 and maximum injection time at 50 ms. Multiply charged ions (z = 2 – 7) with intensity above 10,000 counts were fragmented in HCD (higher collision dissociation) cell at 30% collision energy, and the isolation window at 1.6 Th. The fragment ions were detected in ion trap with AGC at 10,000 and 35 ms or 50 ms maximum injection time. The dynamic exclusion time was set at 45 s and 30 s with ±10 ppm.

### Mass spectrometry data analysis

Raw mass spectrometry data files were analysed with Proteome Discoverer 1.4 (Thermo). Database searches were carried out using Mascot (version 2.2) against the Uniprot human or mouse reference databases (January 2018) appended with the cRAP database (www.thegpm.org/crap/) with the following parameters: Trypsin was set as digestion mode with a maximum of two missed cleavages allowed. Precursor mass tolerance was set to 10 ppm, and fragment mass tolerance set to 0.5 Da. Acetylation at the N-terminus, oxidation of methionine, carbamidomethylation of cysteine, and deamidation of asparagine and glutamine were set as variable modifications. Peptide identifications were set at 1% FDR using Mascot Percolator. Protein identification required at least one peptide with a minimum score of 20. Protein lists from bait and control experiments were analysed with the SAINTexpress (Significance Analysis of INTeractome) algorithm (Teo et al. 2014) to discriminate true interacting proteins from background binders. Using this method, we identified 693 and 1815 unique protein hits from the immunoprecipitation using non-crosslinked and crosslinked cells, respectively. Hits with the highest fold enrichment scores and SAINT probability score cut-off of 1 were members of either the condensin II or SWI/SNF complexes, and are shown in Table 1 and 2 for crosslinked and non-crosslinked cells, respectively.

Identified peptides were analysed using the Contaminant Repository for Affinity Purification at crapome.org to calculate SAINT probability and fold enrichment scores (Mellacheruvu et al. 2013). Top hits were members of either the condensin II or SWI/SNF complex, and are shown in Table 1 and 2, for crosslinked and non-crosslinked samples, respectively.

### Western Blotting

To prepare western blots shown in Figure 4, samples of nuclear lysate and IP were mixed with 4x NuPAGE LDS Sample Buffer (Invitrogen) and run on a 4%–12% Bis-Tris NuPAGE gel (Invitrogen) against a Color Protein Standard broad range ladder (NEB) in NuPAGE MES SDS buffer. Protein was transferred on to a nitrocellulose membrane (Amersham), blocked with 5% milk powder in TBS-T, probed with either primary mouse anti-Smarcc1 (sc-32763) or rabbit anti-NcapH2 (A302-276A, lot 1) and secondary ECL anti-mouse or rabbit IgG HRP-linked (Cytiva). Western was developed on Hyperfilm ECL (Amersham) using ECL substrate (Pierce).

## Acknowledgements

The authors are grateful to Ken-ichi Noma for helpful comments during the preparation of this manuscript.

